# Two pectate lyases from *Caldicellulosiruptor bescii* with the same CALG domain had distinct properties on plant biomass degradation

**DOI:** 10.1101/2020.01.16.910000

**Authors:** Hamed I. Hamouda, Nasir Ali, Hang Su, Jie Feng, Ming Lu, Fu-Li Li

## Abstract

Pectin deconstruction is the initial step in breaking the recalcitrance of plant biomass by using selected microorganisms that carry pectinolytic enzymes. Pectate lyases that cleave α-1,4-galacturonosidic linkage of pectin are widely used in industries, such as paper making and fruit softening. However, reports on pectate lyases with high thermostability are few. Two pectate lyases (*Cb*PL3 and *Cb*PL9) from a thermophilic bacterium *Caldicellulosiruptor bescii* were investigated. Although these two enzymes belonged to different families of polysaccharide lyase, both were Ca^2+^-dependent. Similar biochemical properties were shown under optimized conditions 80 °C–85 °C and pH 8–9. However, the degradation products on pectin and polygalacturonic acids (pGA) were different, revealing the distinct mode of action. A concanavalin A-like lectin/glucanase (CALG) domain, located in the *N*-terminus of two *Cb*PLs, shares 100% amino acid identity. CALG-truncated mutant of *Cb*PL9 showed lower activities than the wild-type, whereas the *Cb*PL3 with CALG knock-out portion was reported with enhanced activities, thereby revealing the different roles of CALG in two *Cb*PLs. I-TASSER predicted that the CALG in two *Cb*PLs is structurally close to the family 66 carbohydrate binding module (CBM66). Furthermore, substrate-binding assay indicated that the catalytic domains in two *Cb*PLs had strong affinities on pectate-related substrates, but CALG showed weak interaction with a number of lignocellulosic carbohydrates, except sodium carboxymethyl cellulose and sodium alginate. Finally, scanning electron microscope analysis and total reducing sugar assay showed that the two enzymes could improve the saccharification of switchgrass. The two *Cb*PLs are impressive sources for degradation of plant biomass.

**Importance:** Thermophilic proteins could be implemented in diverse industrial applications. We sought to characterize two pectate lyases, *Cb*PL3 and *Cb*PL9, from a thermophilic bacterium *Caldicellulosiruptor bescii*. The two enzymes had high optimum temperature, low optimum pH, and good thermostability at evaluated temperature. A family-66 carbohydrate binding module (CBM66) was identified in two *Cb*PLs with sharing 100% amino acid identity. Deletion of CBM66 obviously decreased the activity of *Cb*PL9, but increase the activity and thermostability of *Cb*PL3, suggesting the different roles of CBM66 in two enzymes. Moreover, the degradation products by two *Cb*PLs were different. These results revealed these enzymes could represent a potential pectate lyase for applications in paper and textile industries.

## 1. Introduction

Pectin is an intercellular cement that gives plants structural rigidity; it was discovered at concentrations of 15% to 30% in fruit fibers, vegetables, legumes, and nuts (1). In terms of chemistry, it is an intricate polysaccharide with a molecular weight in the range 20–400 kDa (2). The simplest structure of pectin is a linear polymer of galacturonic acids with α-1,4-galacturonosidic linkages, which is named as homogalacturonan (HG). More complicated forms of pectin are linked by D-galacturonate and L-rhamnose residues, which are designated as types I and II rhamnogalacturonan (RGI and RGII, respectively). The side chains of these RGI and RGII are mostly modified by neutral sugars, such as D-galactose, L-arabinose, D-xylose, and L-fucose (3).

Pectin depolymerization is an initiating step for the bioconversion of plant biomass. Therefore, pectate lyases, a group of enzymes that break down pectin in nature, are used as an additive in generating commercial cellulase cocktails and consolidated-bioprocessing (CBP) microorganisms (4). The mechanism of action of pectate lyase is to cleave the α-1,4-galacturonosidic linkage of polygalacturonic acid (pGA) by β-elimination reaction, thereby forming a double bond between positions C4 and C5 in oligo-galacturonates (1, 4). Through the carbohydrate-active enzyme (CAZy, www.cazy.org) database (5), pectate lyases are classified into families-1, −2, −3, −9, and −10 of polysaccharide lyases. Most of these lyases are identified from microorganisms, such as those from the genera of *Bacillus, Clostridium, Erwinia, Fusarium, Pseudomonas, Streptomyces*, and *Thermoanaerobacter*, which are mostly alkaline (pH 8–11) and Ca^2+^-dependent (4).

Pectate lyases are widely applied in industries, such as paper making, fruit softening, juice and wine clarification, plant fiber processing, and oil extraction (6). Biotechnological usage is highly dependent on the biochemical properties of these enzymes, such as temperature, pH, and salt concentration (1). Commercial pectate lyases were mostly isolated from the mesophilic bacterium *Bacillus*, including *B. subtilis, B. licheniformes, B. cereus, B. circulens, B. pasteurii, B. amyloliquefaciens*, and *B. pumulis* (1, 4). Among them, the recombinant BacPelA from *B. clausii* had high cleavage activity on methylated pectin. Its maximum activities are observed at pH 10.5 and 70 °C and showed the highest degumming efficiency reported to date (7). A pectate lyase pelS6 from *B. amyloliquefaciens* S6 strain displayed highly thermostability and survival in a wide range of pH, which could be suitable for juice clarity (8). To identify thermostable and highly active enzymes, the number of studies investigating thermophilic microorganisms, such as those from the genera of *Thermotoga* and *Caldicellulosiruptor*, has been growing recently (9-14).

A previous study indicated that a pectin-deconstruction gene cluster is vital for the growth of thermophilic *C. bescii* on plant biomass (15). Fig. 1A shows three adjacent genes, *Cbes_1853, Cbes_1854*, and *Cbes_1855* that encode a rhammogalacturonan lyase *Cb*PL11, a pectate lyase *Cb*PL3, and a pectate disaccharide lyase *Cb*PL9, respectively (16). Alahuhta *et al.* presented a crystal structure of the catalytic domain of *Cb*PL3 and described *Cb*PL3 as a unique pectate lyase with a low optimum pH (11, 17). However, the detailed properties of these pectate lyases remained unclear. Meanwhile, *Cb*PL3 and *Cb*PL9 share the same *N*-terminal domain, which is predicted as a concanavalin A-like lectin/glucanase (CALG) domain with 100% protein sequence similarity. The function of this CALG is still unknown. In this study, the pectate lyases *Cb*PL3 and *Cb*PL9 and their truncated mutants were heterologously expressed in *E. coli*. The properties of the different domains in the two enzymes were biochemically and functionally analyzed.

**Fig. 1.**
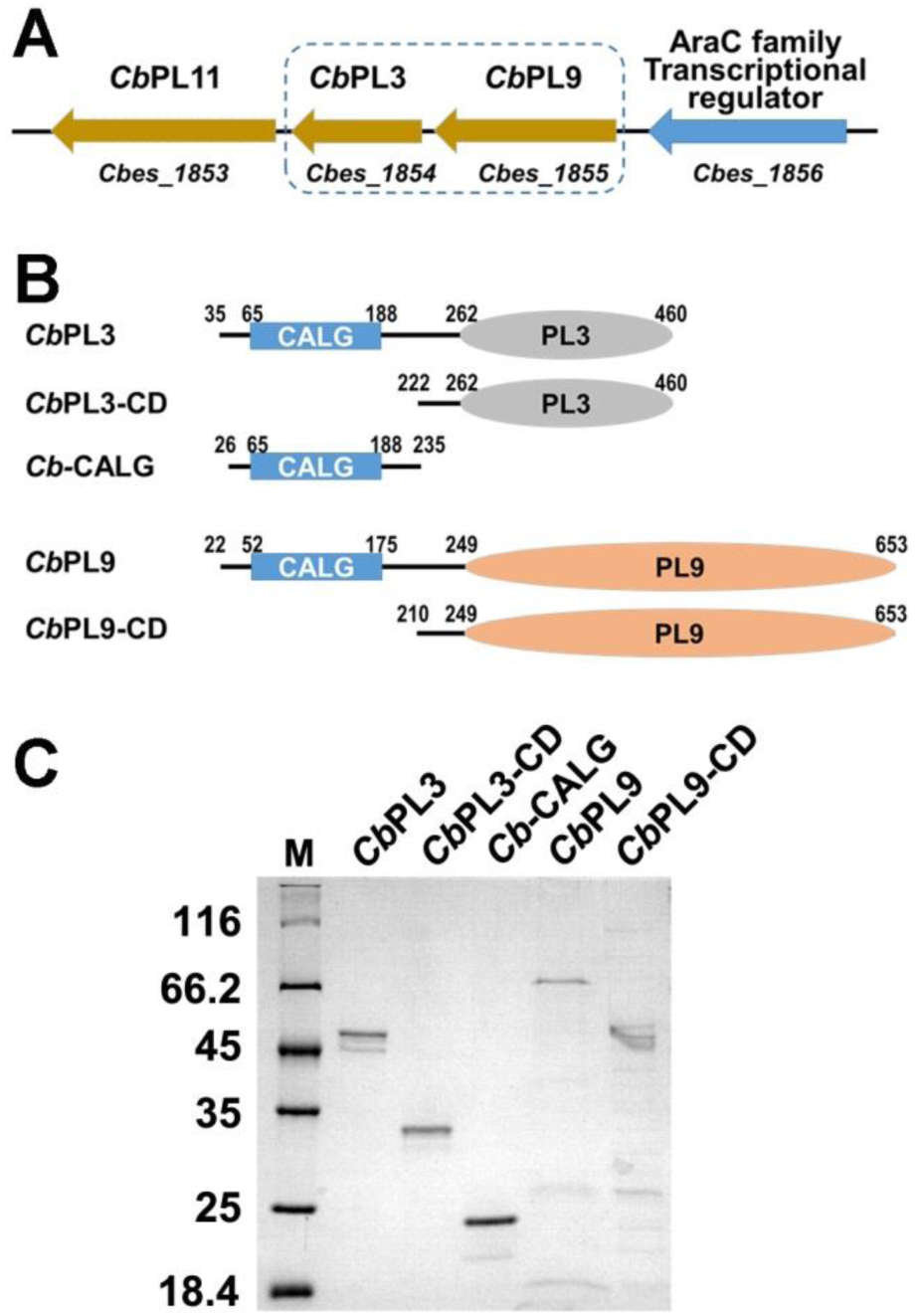
**A.** Gene organization of three pectate lyase genes in a gene cluster of *Caldicellulosiruptor bescii*. **B.** Schematic structures of *Cb*PL3, *Cb*PL9 and their truncated mutants *Cb*PL3-CD, *Cb*PL8-CD and concanavalin A-like lectin/glucanase (CALG). **C.** 12% SDS-PAGE analysis of purified proteins, lane M is protein marker ranged with 18.4-116 kDa.

## 2. Materials and methods

### 2.1 Chemicals and Strains

Mono-galacturonic acid, di-galacturonic acid, tri-galacturonic acid, polygalacturonic acid, citrus pectin, apple pectin, rhamnogalacturonic acid, CMC-Na, Avicel, beechwood xylan, alginate, and cellobiose were purchased from Solarbio Science & Technology (Beijing, China). The vector pEAZY blunt-E2, competent cells *E. coli* BL21 (DE3) pLysS, and Plasmid mini Prep Kit were purchased from TransGen Biotech (Beijing, China). Isopropyl β-D-1-thiogalactopyranoside (IPTG) and ampicillin were purchased from Sigma-Aldrich (St. Louis, MO, USA).

### 2.2 Gene cloning, protein expression, and purification

The genomic DNA was isolated from *C. bescii* (18). Two pectate lyases *Cb*PL3 and *Cb*PL9 were submitted to the NCBI database with GenBank Accession Nos. ACM60942 and ACM60943, respectively (19). The genes of two pectate lyases and their truncated mutants were amplified by polymerase chain reaction and then cloned into pEASY blunt-E2 vector. The primers used in this study are shown in Table 1. The recombinant *E. coli* strains expressing the target proteins were cultivated in a 2 L flask containing 500 mL LB medium mixed with 100 μg/mL ampicillin at 37 °C and agitated at 200 rpm. When OD_600_ of the culture medium reached 0.6–0.8, 0.5 mM IPTG was added to the culture to induce protein expression. The cells were incubated with overnight shaking at 150 rpm at 16 °C. The cell pellets were harvested by centrifugation (8,000×g 10 min, 4 °C) and then dissolved in buffer A (20 mM Tris-HCl, and 300 mM NaCl, pH 8.0). The cell lysates were heated for 15 min at 65 °C and then centrifugation (12,000×g, 30 min, 4 °C) to remove cell debris and denatured proteins after ultra-sonication.

**Table 1.**
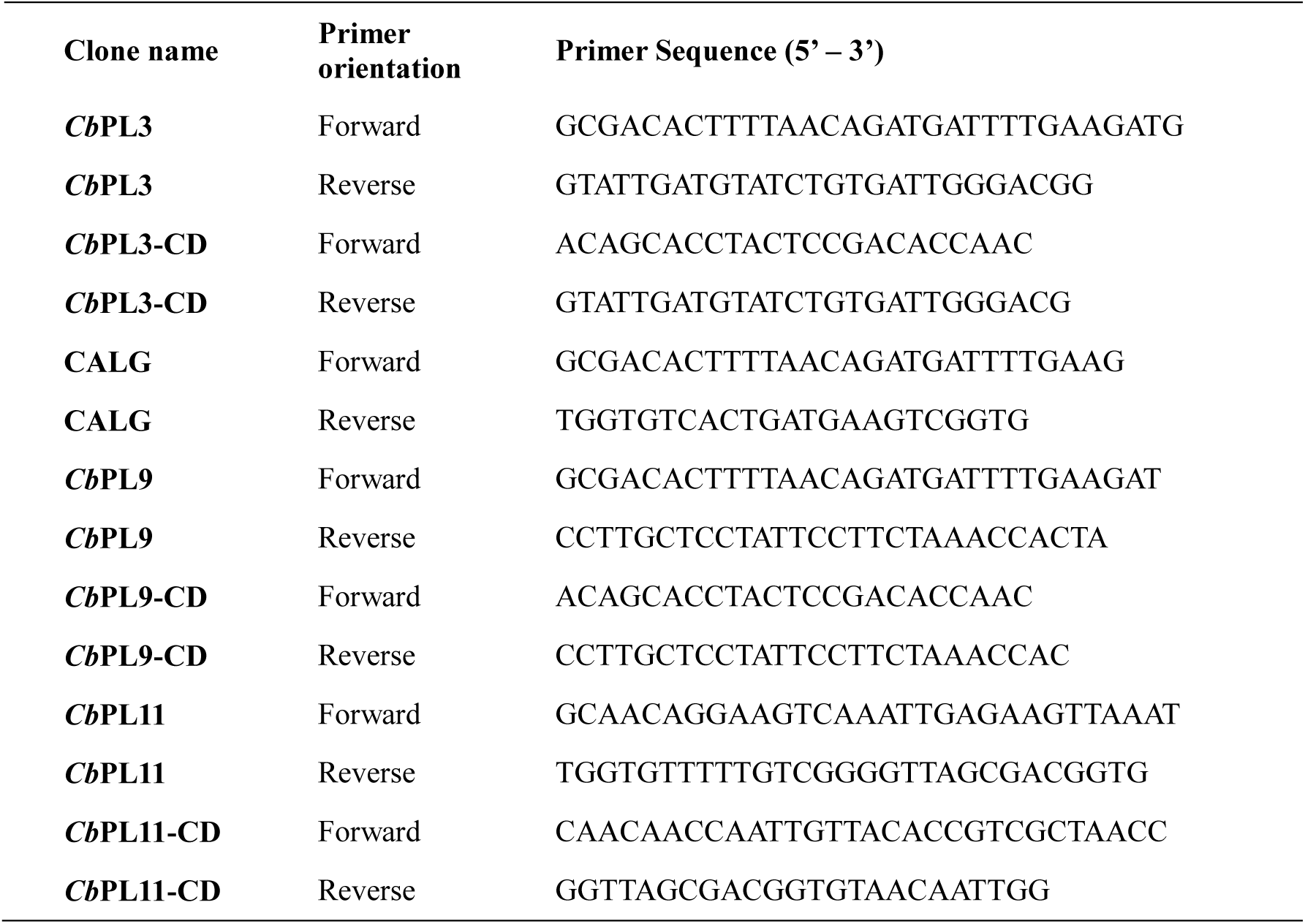
List of used primers.

**Table 2.** Activities of two CbPLs and their catalytic domain on different carbohydrate substrates

| Enzymes | pGA | Citrus<br>pectin | Apple<br>Pectin | Rhamno-GA | CMC-Na | Avicel | Beechwood<br>xylan | Alginate | cellobiose | Inulin<sup>301</sup> |
| --- | --- | --- | --- | --- | --- | --- | --- | --- | --- | --- |
| *CbPL3* | 457.7±4.9 | 231.3 | 15.7 | 7.9 | N.D | N.D | N.D | N.D | N.D | N.D<sup>302</sup> |
| *CbPL3-CD* | 439.2±10.7 | 241.5 | 12.9 | 2.6 | N.D | N.D | N.D | N.D | N.D | N.D<sup>303</sup> |
| *CbPL9* | 21.1±1.1 | 10.6 | 9.9 | 0.7 | N.D | N.D | N.D | N.D | N.D | N.D |
| *CbPL9-CD* | 5.2±0.2 | 1.8 | 1.7 | 0.1 | N.D | N.D | N.D | N.D | N.D | N.D<sup>305</sup> |

The recombinant proteins with 6×His tag were purified using nickel nitrilotriacetic acid (Ni-NTA)-sefinose resin (Sangon, China) in open columns. The open column was equilibrated with 100 ml buffer A (20 mM Tris-HCl, 300 mM NaCl, pH 8.0) and then incubated with supernatant for 20 min. After washing with 100 ml buffer B (buffer A with 40 mM imidazole), the target proteins were eluted with buffer C (buffer A with 300 mM imidazole). The eluted proteins were concentrated using Amicon Ultra filters (Millipore, Ireland), and confirmed by SDS-PAGE.

### 2.3 Activity assay of pectate lyases

The activities of pectate lyases and their catalytic modules were assayed against 0.1% (w/v) of pGA and apple and citrus pectins at appropriate temperatures, pH, and CaCl_2_ concentrations. The absorbances at 235 nm (A_235_) were measured due to the 4,5-unsaturated oligosaccharides. The molar extinction coefficient is 4600 M^-1^cm^-1^ (20). One unit (U) of enzyme activity is the amount of enzyme that produces 1 µmol of 4,5 -unsaturated oligosaccharides per minute. Each experiment was performed in triplicate.

### 2.4 Biochemical characterization of pectate lyases

With 0.1% (w/v) pGA as a substrate, the effect of Ca^2+^ on pectate lyase activity was measured at pH 8.5 and 65 °C by incubating the purified enzyme solutions in 100 mM Tris-Glycine buffer at a Ca^2+^ concentration range of 0–2 mM under standard test conditions.

The effect of pH on pectate lyase activity was measured using 0.1% (w/v) pGA as a substrate at 65 °C and 0.25 mM Ca ^2+^ by incubating the purified enzyme solutions in different buffers, such as sodium citrate buffer (pH 4.0-6.0), Tris-HCl buffer (pH 6.0-9.0), and Glycine-NaOH buffer (pH 9.0-11.0) under standard test conditions.

Furthermore, the effects of temperature on pectate lyase activity were investigated at 55 °C, 65 °C, 75 °C, and 85 °C using 0.1% (w/v) pGA as a substrate at pH 8.5 and 0.25 mM Ca^2+^ to determine the optimal reaction temperature. The purified enzyme solutions were incubated at the abovementioned different temperatures, and residual activities at different times were measured under the standard assay conditions to evaluate the thermal stability of the enzyme.

In addition, the effects of different metal ions and chemicals on pectate lyase activity were measured using 0.1% (w/v) pGA as a substrate at 65 °C, pH 8.5, and 0.25 mM Ca ^2+^ based on the standard assay method. The relative activities of the enzyme were measured with the following metal ions: K^+^, Mn^2+^, Ni^2+^, Hg^2+^, Zn^2+^, Fe^2+^, Fe^3+^, Co^2+^, Cu^2+^, Cd^2+^, Mg^2+^, and Na^+^ at 1 and 5 mM concentrations. EDTA, SDS, Urea, and Guanidine hydrochloride were used at 1% and 5% concentrations.

### 2.5 Analysis of degradation products by *Cb*PLs and their mutants

Degradation of polygalacturonic acid and pectin by pectate lyases was analyzed by TLC by using a solvent system of ethyl acetate: acetic acid: water (4:2:3, v/v/v). The reaction products were visualized by heating the TLC plate at 100 °C for 10 min after spraying with 10% (v/v) sulfuric acid in ethanol. Unsaturated saccharides and standard samples on TLC plates were detected by thiobarbituric acid and o-phenylenediamine staining methods, respectively. Galacturonic acid (M), Di-galacturonic acid (D), and Tri-galacturonic acid (T) (Santa Cruz Biotechnology, Dallas, USA) were used as standards (21).

### 2.6 Scanning electron microscopy (SEM)

Scanning electron microscopy (SEM) was investigated to observe the microstructure and surface morphology of the untreated and enzyme-treated switchgrass leaves. The samples were coated with a 200-Å gold layer using a vacuum sputter, and photos were shot for evidence with SEM (JSM6510LV, Japan) at 10 kV acceleration voltages (22).

## 3. Results and discussion

### 3.1. Cloning and expression of pectate lyases and their truncated mutants from *C. bescii*

A rhammogalacturonan lyase (*Cb*PL11), a pectate lyase (*Cb*PL3), and a pectate disaccharide lyase (*Cb*PL9) were identified in a pectin-depolymerization gene cluster of *C. bescii* (Fig. 1A)(15-17). To determine their roles in pectin deconstruction, we expressed these three lyases and their truncated mutants in *E. coli* protein expression system. Unfortunately, the recombinant forms of *Cb*PL11 and its catalytic module were hardly overexpressed in BL21(DE3) and C41(DE3) due to unknown proteolysis degradation (data not shown). This enzyme contains a catalytic domain that belonged to family-11 polysaccharide lyase (PL) and a family-3 carbohydrate-binding module (CBM3). A previous study demonstrated the lack of glycosylation between two linker modules, which may cause unexpected proteolysis after the heterologous expression of *C. bescii*’s CelA in *B. subtilis* (23, 24). Another two lyases, *Cb*PL3 and *Cb*PL9, contained the same region with 100% protein similarity; this region was predicted to be a concanavalin A-like lectin/glucanase (CALG) domain by NCBI database and Interpro server (25) (Fig. 1B). Phylogenetic analysis suggested that *Cb*PL3 and *Cb*PL9 are more closely related to the pectate lyases from the genus of *Bacillus* than to the ones from other bacteria (Fig. S1). To understand their functions, full-length *Cb*PL3 and *Cb*PL9, and their truncated mutants *Cb*PL3-CD, *Cb*PL9-CD, and CALG were expressed in *E. coli* as confirmed by SDS-PAGE (Fig. 1C).

### 3.2. Biochemical characterization

Pectate lyase activities are mostly Ca^2+^ dependent (4). The crystal structure of *Cb*PL3-CD and tri-galacturonic acid complex reported by Alahuhta M. *et al.* revealed that three Ca^2+^ ions were directly involved in substrate binding (11). To determine the relationship of enzyme activity and Ca^2+^, purified proteins *Cb*PL3, *Cb*PL3-CD, *Cb*PL9, and *Cb*PL9-CD were *in situ* assay in the presence of Ca^2+^ and chelating agent EDTA. As shown in Fig. 2A and S2, the absorption lines at 235 nm were flat (0–30 s) in the absence of Ca^2+^. The absorption values increased (70–100 s) after dropping the Ca^2+^ ions (0.1 mM), which were quenched by adding 0.5 mM EDTA (130–160 s). Finally, the absorption values were increased by adding Ca^2+^ (1 mM) (190–220 s). Furthermore, relative activities were examined in the presence of different concentrations of Ca^2+^ ions (Fig. 2B). Two full-length pectate lyases and their catalytic modules had their highest lyase activities at 0.25 mM Ca^2+^, except for *Cb*PL9 at 0.125 mM Ca^2+^ (Fig. 2B). Pectate lyases *Cb*PL3 and *Cb*PL9 are Ca^2+^ dependent. The binding affinity of Ca^2+^ with the catalytic regions of these two pectate lyases was weak in the absence of substrate, and detecting the lyase activities of these purified proteins from *E. coli* was not possible in the absence of Ca^2+^.

**Fig. 2.**
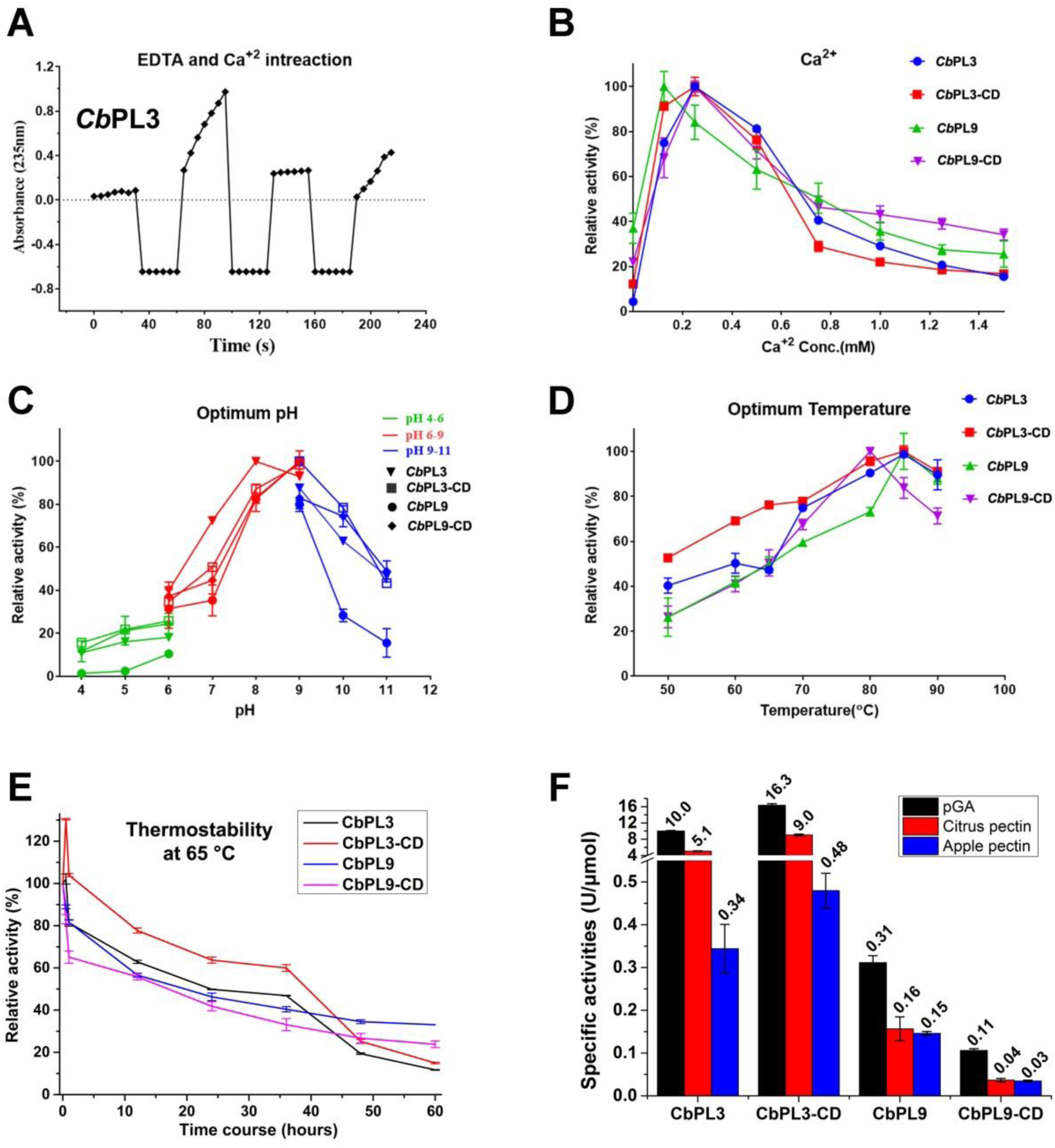
Enzyme characterization: **(A)** Interaction of calcium and chelating agent with *Cb*PL3. **(B)** Optimum Calcium concentration, **(C)** Optimum pHs, **(D)** Optimum temperatures. **(E)** Thermostabilities at 65 °C of *Cb*PL3, *C*bPL9, *Cb*PL3-CD and *Cb*PL9-CD. **(F)** Specific activities (U/μmol) of pectate lyases and their truncated mutants with synthetic substrate (pGA) and natural substrates (Citrus pectin and Apple pectin).

Moreover, the effect of metal ions was examined using pectate lyases and their catalytic modules in the presence of 0.25 mM Ca^2+^. The activities were strongly inhibited by most of the divalent cations (Tables S1 and S2). Optimum pH assay revealed that the enzymes *Cb*PL3-CD, *Cb*PL9, and *Cb*PL9-CD exhibited their best activities at pH 9.0, as shown in Fig. 2C. The wild-type *Cb*PL3 exhibited its optimal activity at pH 8.5 and retained over 10% of activity at pH 7 (11). The data shown in Fig. 2C verified that *Cb*PL3 had a broad pH range, and over 75% of activity was retained at pH 7–10. This phenomenon was possibly due to the difference in the two buffer recipes. Regarding the optimum temperature, the enzymes *Cb*PL3, *Cb*PL3-CD, and *Cb*PL9 exhibited their highest activities at 85 °C, whereas *Cb*PL9-CD showed activity at 80 °C, as shown in Fig. 2D. The half-lives of these enzymes were more than 15 h at 65°C, thereby suggesting better thermal stability at elevated temperatures (Figs. 2E and S2). Finally, specific activities (U/μmol) of these enzymes were examined under optimum conditions of 85 °C, pH 9, and 0.25 mM Ca ^2+^ using various substrates, as shown in Fig. 2F. *Cb*PL3-CD showed higher activities than *Cb*PL3, whereas *Cb*PL9-CD activity was reportedly lower than *Cb*PL9’s, thereby suggesting the different effects of CALG domain on the two pectate lyases. Kinetic analysis of these enzymes demonstrated that the deletion of CALG domain reduced the catalytic efficiency of the two *Cb*PLs by decreasing their turnover rates (*k*cat) (Table S3). Overall, knockout of CALG domain did not significantly influence the two *Cb*PLs but noticeably affected *Cb*PL9 by lowering its activity.

### 3.3. Modes of action

To understand the modes of action, the *Cb*PLs and their truncated mutants were investigated with a substrate-binding assay. The full-length *Cb*PLs and its truncated mutants showed appropriate shifts in the absence of substrates in native protein gels (arrows in Fig. 3A). The protein bands were showed in loading wells in the presence of soluble pGA or citrus pectin, thereby suggesting the formation of the *Cb*PLs-soluble substrates complexes. However, CALG showed the similar protein band positions in the absence and presence of soluble substrates. The catalytic module in the two *Cb*PLs but not CALG domain has strong interaction with soluble pectin-related substrates. We examined the binding capacities of these proteins with an insoluble substrate (switchgrass). These proteins showed their bands in a denatured SDS-PAGE gel in the absence of switchgrass (Fig. 3B). The proteins, except CALG, totally disappeared in the supernatant after centrifugation with insoluble switchgrass. Catalytic modules of *Cb*PL3 and *Cb*PL9 had strong interaction with the insoluble switchgrass, whereas CALG showed a moderate interaction.

**Fig. 3.**
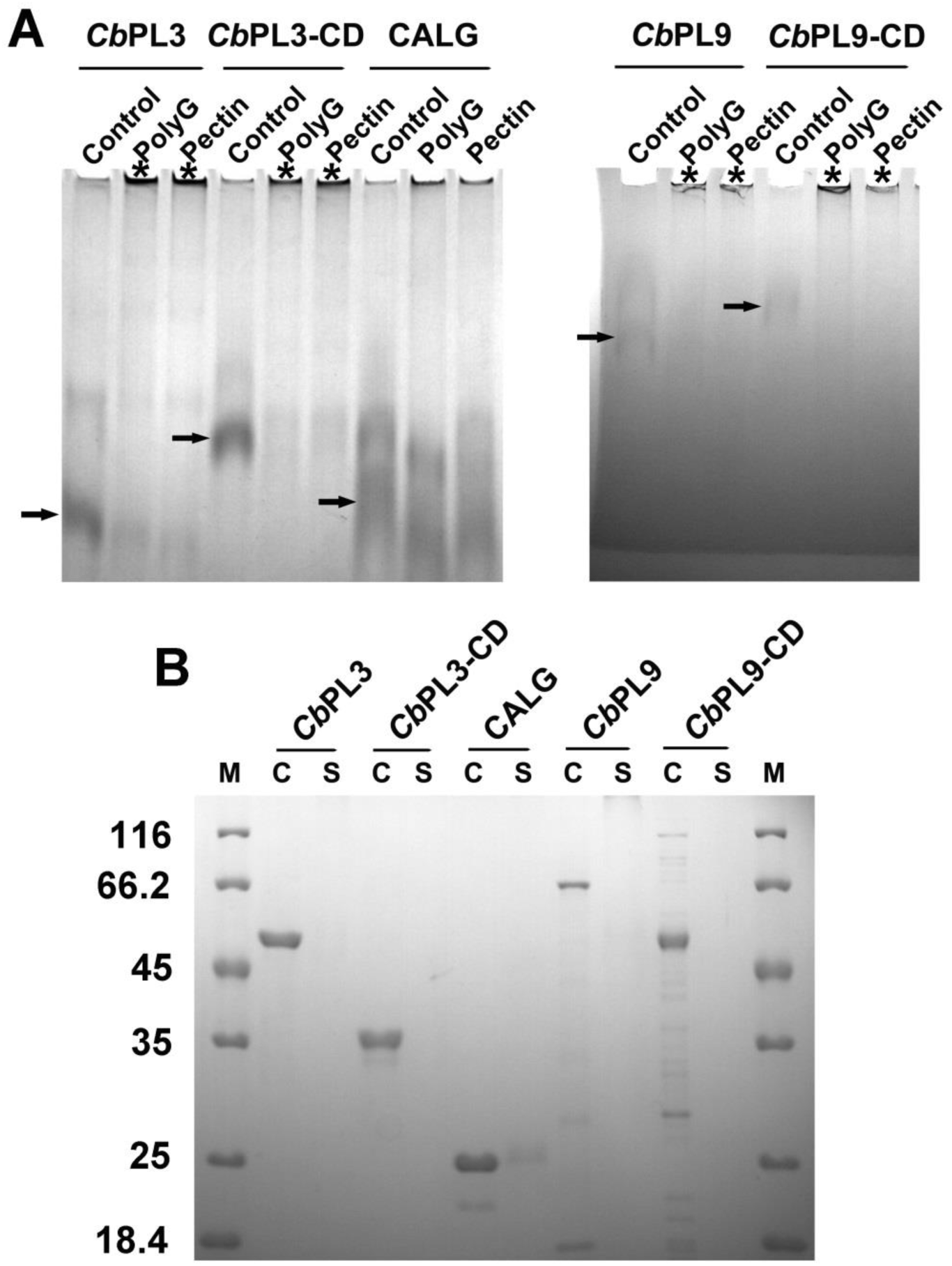
Carbohydrate-substrate binding assay. **(A)** The target proteins were loaded in 8% native gel in the absence or presence of soluble substrates, such as pGA and citrus pectin. Control was labelled by arrows and the protein and substrate complex was marked by *. **(B)** The target proteins were loaded in 12% SDS-PAGE before or after incubate with insoluble natural substrate switchgrass.

The degradation products of *Cb*PLs in the presence of pGA were then analyzed by thin layer chromatography (TLC). The main products of *Cb*PL3 on pGA were unsaturated di-, tri, and tetra-galacturonic acids (GAs) in the initial phase of the reaction (Fig. 4A). The end products were confirmed as unsaturated di- and tri-GAs. For *Cb*PL9, the end products were mainly unsaturated tri- and tetra-GAs together with a slight amount of di-GA (Fig. 4B). Therefore, two *Cb*PLs were endo-type pectate lyases with different action modes. Furthermore, deletion of CALG did not change the profiles of the degradation products of *Cb*PL3 and *Cb*PL9 (Fig. 4B).

**Fig. 4.**
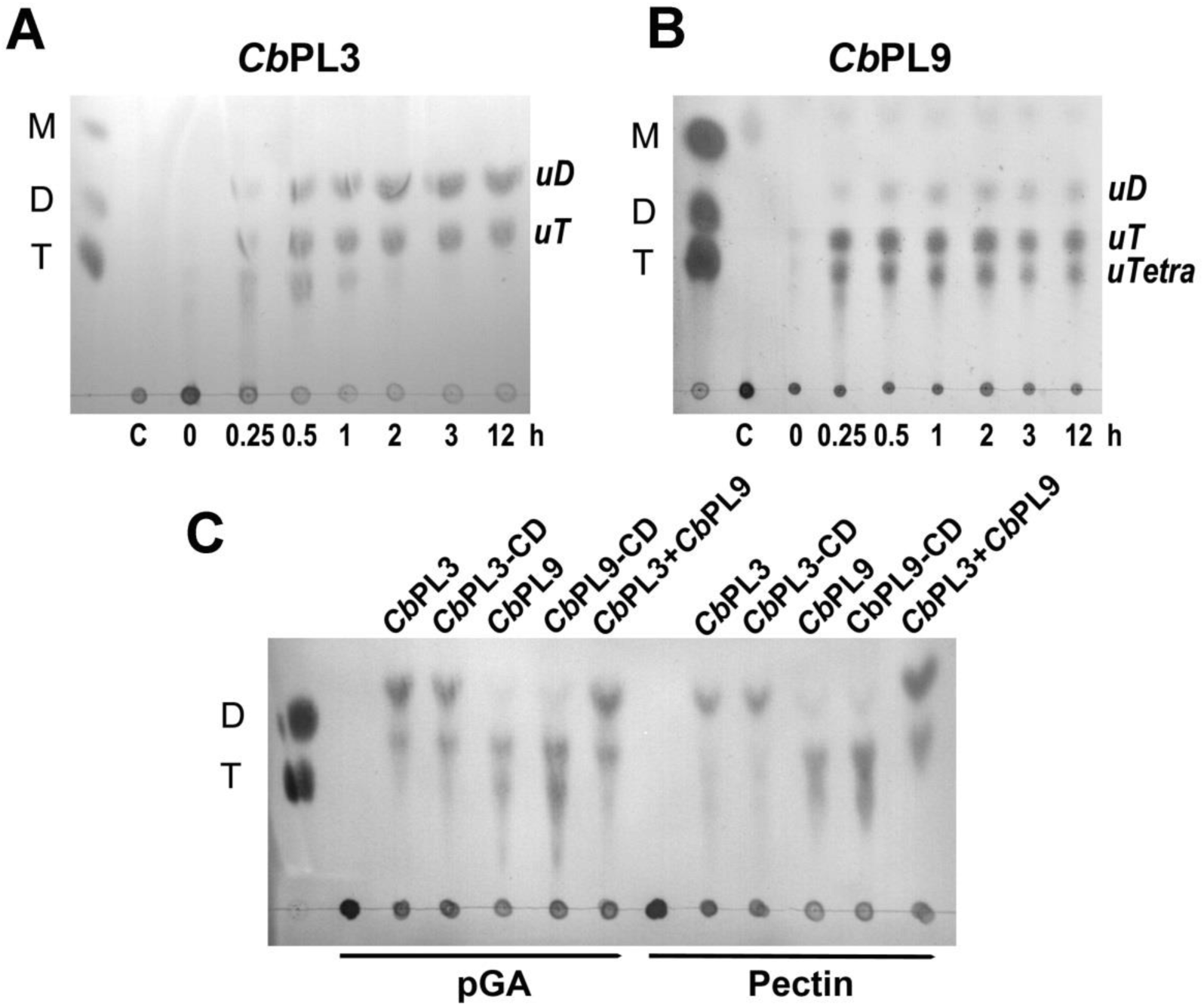
Time-course degradation products of polygalacturonic acid (pGA) by *Cb*PL3 **(A)** and *Cb*PL9 **(B)** by using TLC analysis. Marker contained the standard chemicals such as Mono (M), Di (D) and Tri (T) galacturonic acid; C, pGA was incubated at the absence of enzyme for 12 h, and the following dots show products liberated after 0.0, 0.25, 0.5, 1.0, 2.0, 3.0 and 12.0 h of enzymatic action. **(C)** Degradation products of pGA and pectin by one or combination of different enzymes.

### 3.4. Characterization of CALG

The CALG domain is a protein superfamily with 12–14 anti-parallel β-strands arranged in two curved sheets (26). According to the diverse function, the superfamily CALG was classified into six families, namely, endoglucanase, xylanase, cellobiohydrolase (CBH), β-1,3(−1,4)-glucanase, alginate lyases, and peptidase (26, 27). However, the hydrolysis activities of *Cb*PL3, *Cb*PL9, and CALG were undetectable using the DNS method in the presence of several carbohydrate substrates, such as sodium carboxymethyl cellulose (CMC-Na), beechwood xylan, alginate, cellobiose, and avicel (Table S4). Meanwhile, native protein electrophoresis showed that CALG and the CALG-containing protein, such as *Cb*PL3, has strong substrate-binding affinities with the CMC-Na and alginate, but not with avicel, beechwood xylan, and cellobiose (Figs. 5A and S4). Moreover, CALG binds weakly with pectin and pGA (Fig. 3A) but strongly with intricate plant biomass, such as switchgrass (Fig. 3B). Therefore, CALG acts as a binding module rather than a catalytic module. Using a structure and function predication server, I-TASSER (28), CALG was predicted to be structurally close to the family-66 carbohydrate binding module from *B. subtilis* (*Bs*CBM66) (PDB No. 4AZZ) (Fig. 5B), which was located in the C-terminus of an exo-acting β-fructosidase (29). The substrate binding sites in CALG are different from *Bs*CBM66 (Fig. 5C). Therefore, neither β-fructosidase activity nor binding affinity of CALG and *Cb*PL3 with inulin was detected (Table S4 and Figs. 5A and S4). Overall, CALG in two *Cb*PLs had neither glycoside hydrolase nor polysaccharide lyase function but showed strong carbohydrate binding affinity with CMC-Na, sodium alginate, and switchgrass.

**Fig. 5.**
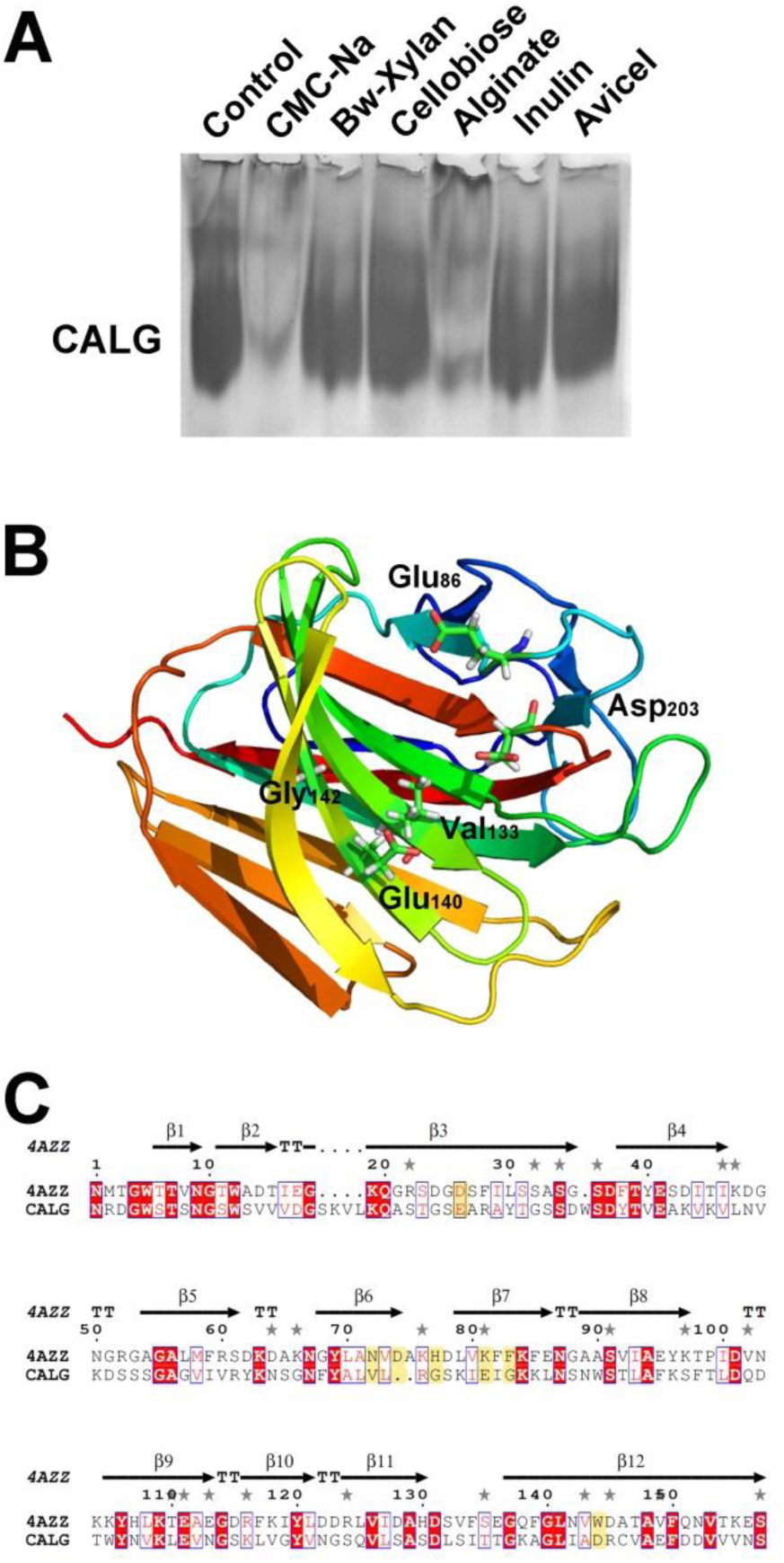
**(A)** Binding assay of concanavalin A-like lectin/glucanase (CALG) domain with different carbohydrate substrates using a native electrophoretic gel. **(B)** The predicated structure and substrate-binding sites of CALG using I-TASSER server. **(C)** Protein alignment of *Bs*CBM66 (PDB: 4AZZ) from *Bacillus subtilis* and CALG domain.

### 3.5. SEM and reducing sugar analysis on plant biomass

Morphological changes in switchgrasses (SG) by two *Cb*PLs were investigated using a scanning electron microscope (SEM). As shown in Fig. 6A, the surface of untreated SG leaves was flat and smooth, and no fiber bundles were observed. However, the surface morphologies of *Cb*PL3- and *Cb*PL9-treated SG leaves were significantly changed (Figs. 6B and 6D). The fiber bundles of treated leaves are exposed with many deep longitudinal cracks from the inside (22). The porosity and permeability of the surface increased, thereby enhancing the accessibility of the cellulase mixture to utilize cellulose and hemicellulose. Together with cellulase cocktail, more splits and fractures in the morphology were observed (Figs. 6C and 6E). Meanwhile, a significant increase in the total free sugars of the tested samples were shown compared with the control (Fig. 7), thereby indicating the effect of *Cb*PLs on natural biomass hydrolysis.

**Fig. 6.**
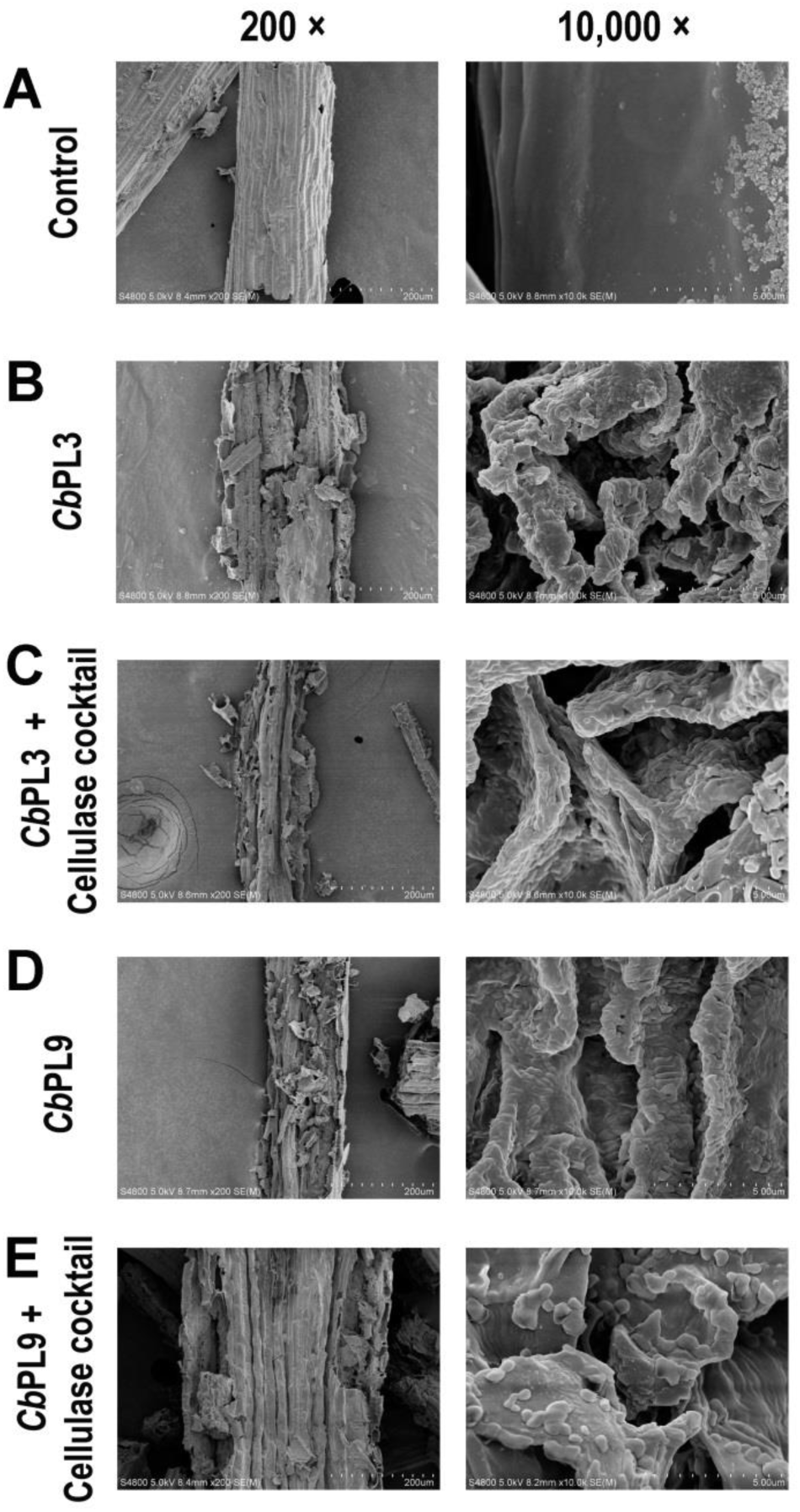
Scanning electron microscope analysis of treated Switch grass at 200 X and 10,000 X, respectively. **(A)** Negative control (with buffer only); **(B)** after *Pectate lyase* (*Cb*PL3); **(B)** after *Cb*PL3 and cellulase cocktail; **(D)** after *Pectate lyase* (*Cb*PL9); **(E)** after *Cb*PL9 and cellulase cocktail.

## Acknowledgements

This work was supported by the National Natural Science Foundation of China (Nos. 31770077 and 31400060) and the “Transformational Technologies for Clean Energy and Demonstration”, Strategic Priority Research Program of the Chinese Academy of Sciences (XDA 21060400). We are also grateful to “CAS-TWAS President’s PhD Fellowship Programme” and “CAS President’s International Fellowship Initiative (PIFI) program” for a part of financial support.

